# A Proof-of-Concept Study on Attached *Spirulina* Cultivation Under Microgravity and Partial Gravity Aboard the ISS

**DOI:** 10.64898/2026.10.27.762075

**Authors:** Takashi Inada, Motoshi Kamada, Hayato Ohkuma, Toru Shimazu, Junya Kirima, Yasuyuki Kikuchi, Haruo Kasahara, Tomokazu Yamazaki, Kana Kuriyama, Eriko Moriyama, Shinichi Aoki, Takanori Hoshino

## Abstract

Microalgal attached cultivation circumvents gas-liquid separation issues of liquid-suspended cultivation, yet performance under reduced gravity remains unevaluated. As a proof-of-concept, we evaluated *Arthrospira platensis* (*Spirulina*) cultivation under artificial 1G, 1/6G, and microgravity (µG) on the International Space Station. *Spirulina* expanded without cell detachment, medium floating, or component separation, with relative radial growth rates of 0.28%^0.5^ day^-1^ (1G), 0.22%^0.5^ day^-1^ (1/6G), and 0.17– 0.21%^0.5^ day^-1^ (µG), demonstrating feasibility in space.

---

Long-duration human space exploration, such as missions to the Moon and Mars, demands sustainable Regenerative Life-Support Systems (RLSS) to minimize resupply requirements from Earth [1, 2]. In closed-loop RLSS architectures, microalgae have been considered one of the most promising biological solutions for O_2_ regeneration, CO_2_ fixation, and food production [3, 4].

Despite this biological potential, traditional liquid-suspended cultivation poses technical challenges in lunar and space environments. For example, liquid-suspended cultivation of *Limnospira indica* on the International Space Station (ISS) using gas-permeable membranes showed operational limits [5]. Without mechanical mixing, liquid boundary layers and bubble accumulation restricted CO_2_ supply and O_2_ removal (Figure 1, left).

**Figure 1.**
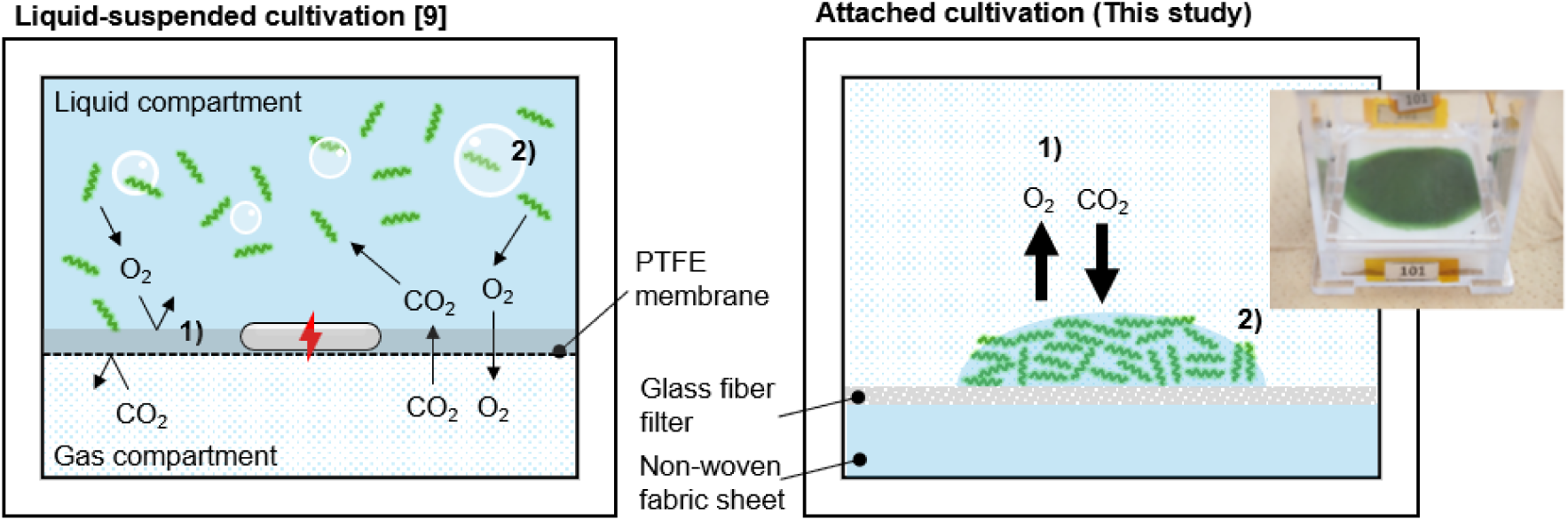
Schematic comparison of microalgal cultivation systems under µG. (Left) Liquid-suspended cultivation in a previous study [9] faces operational issues: 1) a liquid boundary layer, and 2) bubble accumulation. (Right) The proposed attached cultivation eliminates these issues by providing 1) direct gas exchange, and 2) a bubble-free environment.

To circumvent these constraints of liquid-suspended cultivation, we focused on an attached cultivation system. Through high-density cultivation of microalgae on a water-holding substrate saturated with medium, this approach directly exposes cells to the gas phase, facilitating direct gas exchange and creating a bubble-free environment, which intrinsically eliminates the need for gas–liquid separation equipment under reduced gravity. To verify the feasibility of this system as a practical solution, we must confirm that microalgae can grow on the substrate under reduced gravity without critical operational issues. Therefore, the primary objectives of this study were to demonstrate the initial proof-of-concept of an attached *Arthrospira platensis* NIES-39 cultivation system. Specifically, we aimed to confirm (1) the physical stability of the system and (2) expansion performance of *Spirulina* biomass under reduced gravity (Figure 1, right). To evaluate the physical stability of the system, we examined time-lapse images for cell detachment, medium floating, and component separation under artificial 1G, 1/6G, and microgravity (µG). To evaluate the expansion performance of *Spirulina* biomass, we quantified the relative radial growth rates based on green area expansion across three gravity levels (1G, µG, and 1/6G).

We also evaluated the impact of pre-culture cold storage (up to 15 days) on expansion. Time-lapse imaging during experiments on the ISS, captured by the internal camera of the Plant Experiment Unit (PEU) housed within the Cell Biology Experiment Facility (CBEF), provided visual evidence of *Spirulina* expansion behavior on the substrate under different gravity conditions (Figure 2, Supplementary Movies 1–4). In all experimental groups (1G, µG, and 1/6G), the microalgal biomass steadily expanded across the grass fiber filter from Day 0 to Day 8, without any bleaching or decay.

**Figure 2.**
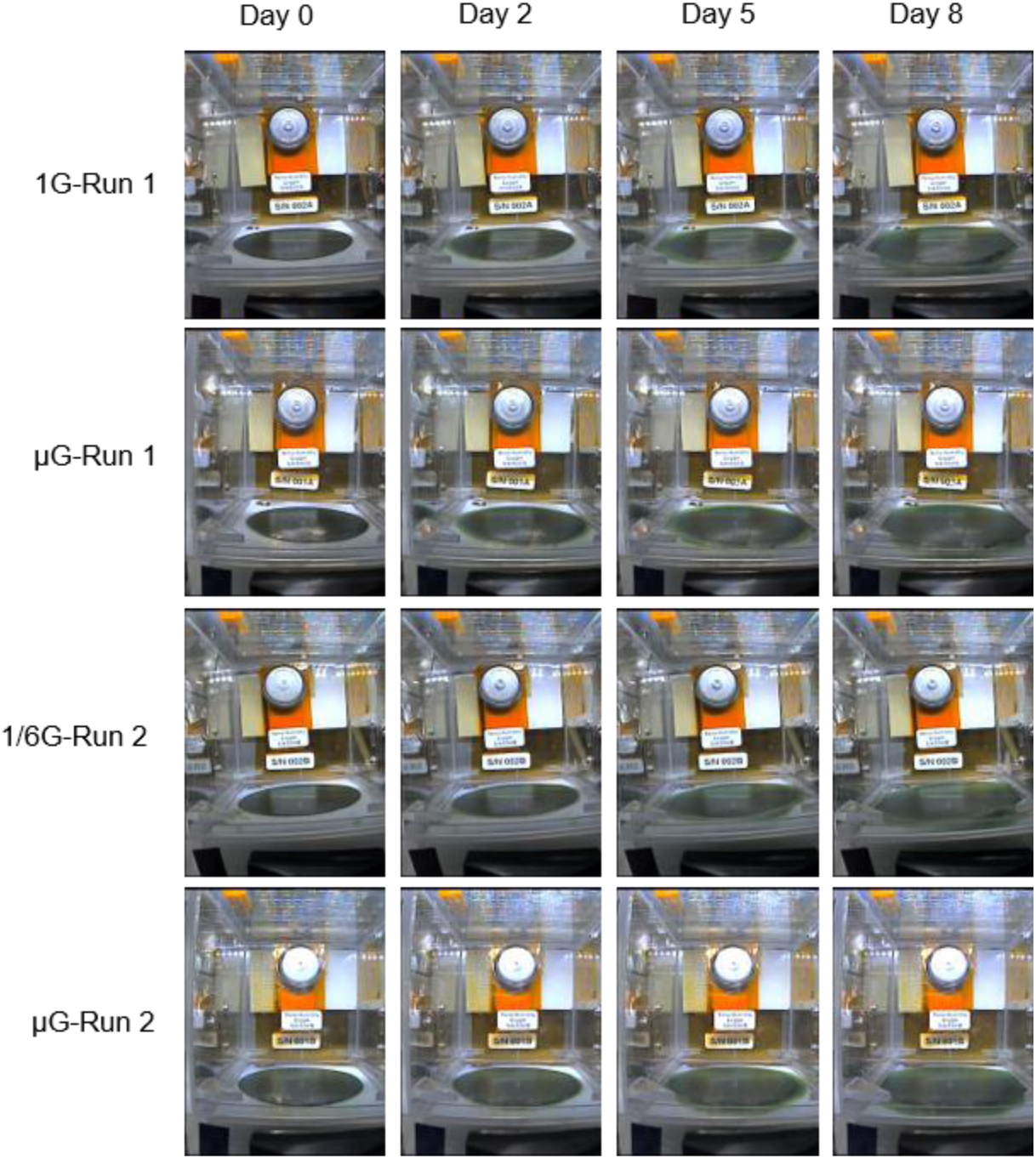
Time-lapse images of attached *Spirulina* cultivation under 1G, µG, and 1/6G, demonstrating continuous biomass expansion without cell detachment, medium floating, or component separation under reduced gravity on the ISS. These images are shown at selected intervals (Days 0, 2, 5, and 8). Each panel represents a horizontal field of view of approximately 40 mm.

Although biomass expansion patterns were generally similar across all gravity levels, we observed marginally reduced expansion in some µG samples in Run 2. From an engineering perspective, time-lapse images not only indicated that no cells detached from the substrate surface but also confirmed no change in the relative positions of the internal components of the vessel, demonstrating that the glass fiber filter and non-woven fabric sheet remained attached without floating or separation under reduced gravity conditions on the ISS. Even with condensed water droplets accumulated on the inner walls of the vessel due to the lack of natural convection under reduced gravity, no liquid floated away or escaped from the non-woven fabric. The non-woven fabric sheet held the Spirulina–Ogawa–Terui (SOT) medium through capillary action, while the glass fiber filter supported *Spirulina* cell attachment with direct gas-phase exposure.

The surface tension between these two layers likely maintained the structure and prevented fluid separation under reduced gravity. These images also indicated that the area under the LED light source remained clear, presumably attributable to thermal dissipation from the LEDs, suggesting that the reduction in irradiance by condensation was minimal. However, during long-term cultivation, accumulated condensation could decrease light intensity. To prevent these issues in future designs, using gentle forced airflow or anti-fog coatings on the inner walls will be needed to manage humidity and light. The uniform surface expansion and stable cell attachment suggest that, in future applications, high-density biomass with low water content of approximately 90 wt% (Supplementary Table 1) could be harvested directly via scraping under reduced gravity, without requiring liquid dewatering equipment.

To quantitatively evaluate the expansion behaviors, the projected Spirulina area was non-destructively extracted from time-lapse images using a semi-automated Fiji image-processing pipeline (Figure 3A, B, Supplementary Movies 1–4; see Methods for details). The relative radial growth rates, calculated from area expansion, varied across different gravity conditions (Figure 3C). In Run 1, the relative radial growth rate under µG (0.21 ± 0.02%^0.5^ day^-1^, *n* = 4) showed a slightly lower mean value than that under 1G (0.28 ± 0.02%^0.5^ day^-1^, *n* = 4), yet no statistically significant difference was detected between the two conditions (*p* = 0.072, Student’s *t*-test).

**Figure 3.**
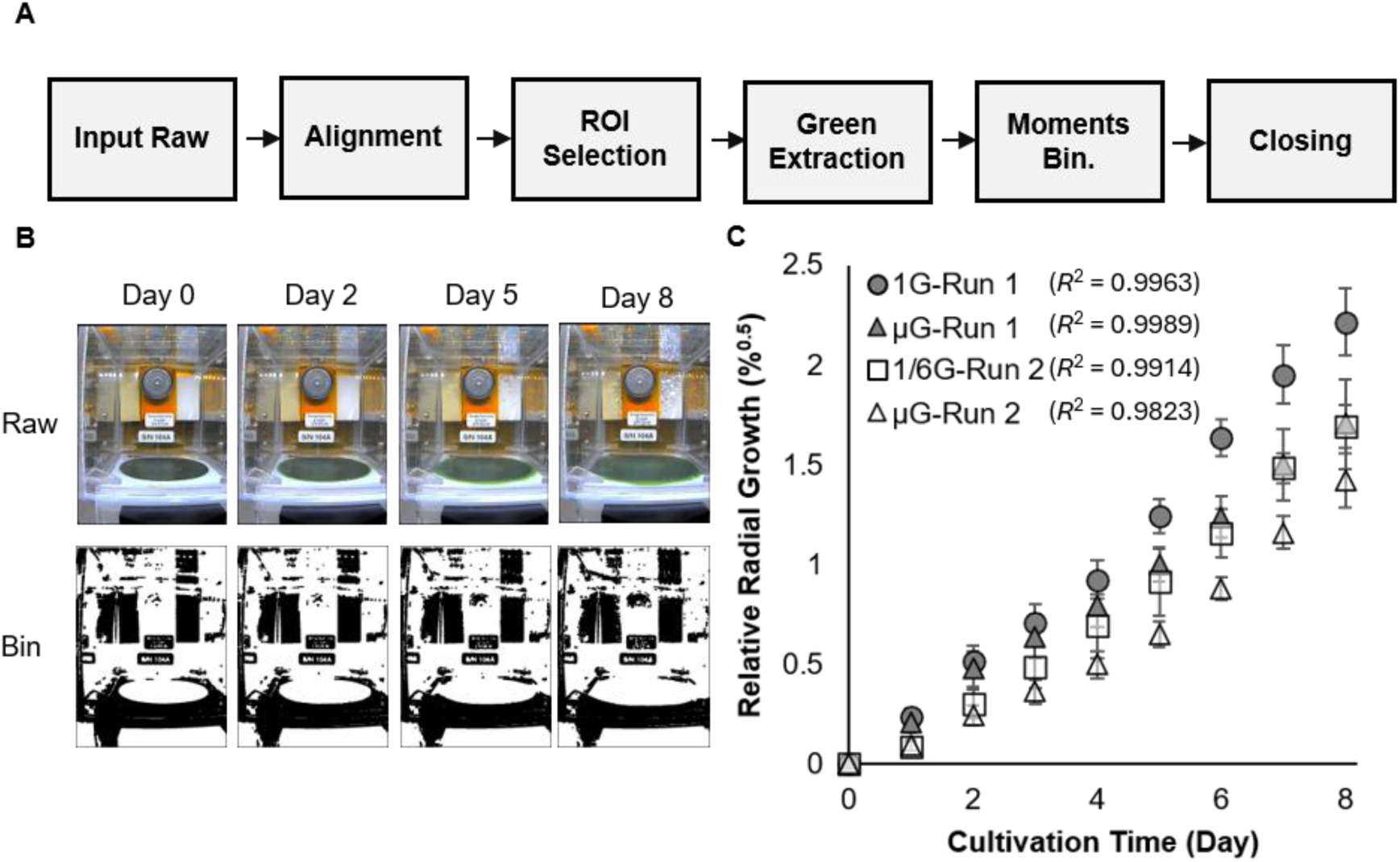
Quantification of Spirulina biomass expansion under 1G, µG, and 1/6G using the developed image-processing pipeline. (A) Workflow of the semi-automated image processing pipeline performed in Fiji, including image alignment, ROI selection, green channel extraction, Moments-based binarization, and morphological closing (two cycles of dilation and erosion). (B) Raw (upper) and binarized (lower) time-lapse images of Spirulina on the supporting materials from Day 0 to Day 8. (C) Relative radial growth curves based on the extracted microalgal area under 1G, µG, and 1/6G conditions. Closed circles, 1G-Run1; closed triangles, µG-Run1; open squares, 1/6G-Run 2; open triangles, µG-Run 2. Data points represent the mean values of four independent replicates (*n* = 4), and error bars indicate the standard error (SE). Coefficients of determination (*R*^2^) from linear regression analysis for each condition are displayed within the plot.

In Run 2, the relative radial growth rate under 1/6G (0.22 ± 0.01%^0.5^ day^-1^, *n* = 4) showed a higher mean value than that under µG (0.17 ± 0.01%^0.5^ day^-1^, *n* = 4), but no significant difference was detected (*p* = 0.053, Student’s *t*-test). Although no statistically significant differences were detected, the data showed a trend towards a lower relative radial growth rate under µG compared to higher gravity (1G or 1/6G). Nevertheless, continuous biomass expansion was maintained across all tested conditions.

The differences in relative radial growth rates across gravity levels likely reflect physical constraints rather than intrinsic physiological responses because cyanobacteria lack gravity-sensing organs and do not exhibit stress responses to gravity changes [6]. Two physical factors can explain the lower radial growth rate observed under µG. First, the lack of natural convection in microgravity can form a stagnant gas boundary layer around the cells, restricting gas exchange of CO_2_ and O_2_ [7]. Second, microgravity can induce a morphological shift toward vertical biofilm thickening rather than horizontal spreading, as observed in other spaceflight microorganisms [8]. Because the internal RGB camera in the Plant Experiment Unit (PEU, JAXA, Tsukuba, Japan) could not quantify biomass thickness, vertical growth could lead to an underestimation of total biomass from 2D area measurements. Future missions incorporating gas exchange sensors and 3D optical imaging will be needed to distinguish between these two mechanisms. Nevertheless, continuous biomass expansion was maintained across all gravity conditions (1G, 1/6G, and µG), confirming the operational feasibility of attached microalgal cultivation in space. Comparing the two microgravity controls, µG in Run 1 exhibited a slightly higher mean area relative radial growth rate than µG in Run 2, although no statistically significant difference was detected (*p* = 0.275, Student’s *t*-test). Preliminary ground evaluation confirmed that cold storage for up to 4 weeks had a negligible effect on *Spirulina* growth, indicating that the cells remained physiologically active (Supplementary Figure 1). Thus, the minor variance between the two µG runs likely reflects small operational differences during manual setup or subtle fluctuations in irradiance rather than biological inhibition.

The entire experiment, including 15 days of pre-culture cold storage and 8 days of monitoring in space, was successfully completed without operational issues associated with liquid-suspended cultivation, such as fluid separation or bubble accumulation.

In conclusion, this study demonstrates both the physical stability of the attached cultivation system and the expansion performance of *Spirulina* biomass under µG and 1/6G aboard the ISS, providing initial proof-of-concept. The continuous expansion of *Spirulina* biomass, supported by capillary forces and surface tension for liquid retention and cell adhesion, shows that this system provides a practical approach for the cultivation of microalgae under reduced gravity. Combined with non-destructive image analysis, these findings offer a useful basis for designing microalgal photobioreactors for future RLSS. While direct measurement of gas exchange efficiency remains a necessary next step, ongoing post-flight analyses of biomass will further evaluate its nutritional quality and genetic stability for long-term space missions.

## Methods

### Microalgal Strain and Pre-cultivation

*Arthrospira platensis* NIES-39 was obtained from the Microbial Culture Collection at the National Institute for Environmental Studies (NIES, Tsukuba, Japan). The SOT medium was prepared according to the formulation designated by NIES [10], containing (per liter): 16.8 g NaHCO_3_, 0.5 g K_2_HPO_4_, 2.5 g NaNO_3_, 1.0 g K_2_SO_4_, 1.0 g NaCl, 0.2 g MgSO_4_·7H_2_O, 0.01 g CaCl_2_·2H_2_O, 0.01 g FeSO_4_·7H_2_O, 0.08 g EDTA, and 1.0 mL of micronutrient solution (containing H_3_BO_4_, MnSO_4_·5H_2_O, ZnSO_4_·7H_2_O, CuSO4·5H_2_O, and Na_2_MoO_4_·4H_2_O). After dissolving ingredients, the SOT medium was filter-sterilized through a 0.22-µm PES membrane filtration unit (Nalgene Rapid-Flow, Thermo Fisher Scientific, Waltham, MA, USA).

*Spirulina* was pre-cultured in a 300 mL Erlenmeyer flask containing 250 mL SOT medium. Cultivation was performed at 25 °C under continuous light at 78 µmol photons m^-2^ s^-1^ using a plant growth LED light (LSB-27K; Takasho Co., Ltd., Wakayama, Japan) with air bubbling for 7 days using an incubator at the NASA Space Station Processing Facility (SSPF) in Kennedy Space Center. Air bubbling was supplied to the culture using a silicone tube (inner diameter: 4 mm, outer diameter: 6 mm; AS ONE Corporation, Osaka, Japan) connected to an air pump (Whisper 60; Spectrum Brands, Middleton, WI, USA) with an inline 0.2-µm PTFE syringe filter (Ekicrodisk E255; Pall Corporation, Port Washington, NY, USA) for sterile air supply. The flow rate was adjusted to approximately one bubble per second.

### Sample Inoculation, Vessel Assembly, and Cold Transportation to the ISS

To determine the dry biomass density, 3–15 mL of *Spirulina* culture was filtered onto a glass fiber filter (Whatman GF/C; GE Healthcare, Chicago, IL, USA), which had been pre-dried overnight at 90 °C and weighed. The retained biomass was rinsed twice with 50 mL ultrapure water to remove residual salts from the medium. The filter containing the biomass was then dried at 90 °C overnight and weighed gravimetrically. Using a calibration curve established from these dry weight measurements, the required volume of *Spirulina* culture was calculated to achieve a target seeding mass of 15 mg.

Prior to assembly, the 40 mm × 50 mm glass fiber filters (GB-100R; ADVANTEC TOYO, Tokyo, Japan) and non-woven fabric sheets (Germguard; Toyobo Co., Ltd., Osaka, Japan) were autoclaved at 121 °C for 20 min. For final flight preparation at SSPF, *Spirulina* inoculum was seeded onto the pre-sterilized glass fiber filter via vacuum filtration as a circular area with a diameter of 38 mm, defined using a custom-fabricated silicone rubber ring (outer diameter: 50 mm, inner diameter: 35 mm, thickness: 0.5 mm, Shore A hardness: 50) placed between the filter and the filtration holder.

For the assembly of the PEU culture vessel, five pieces of the autoclaved nonwoven fabric sheets were saturated with 20 mL of SOT medium and placed at the bottom of a transparent polycarbonate vessel (outer dimensions: 58 mm × 66 mm × 64 mm). The *Spirulina*-seeded glass fiber filter was then placed on top of the saturated sheets.

The assembled culture vessels were stored at 4 °C and transferred to the NASA Cold Stowage team within 24 hours of assembly. The samples were launched aboard the Cygnus NG-24 resupply mission to the ISS on April 11, 2026. Cultivation for the Run 1 experiment began on April 15, and for the Run 2 experiment on April 23. Thus, prior to the start of orbital cultivation, the total 4 °C storage period was 7 days for Run 1 and 15 days for Run 2.

Details of the specific cultivation conditions and experimental setups used for the preliminary study on cold storage (Supplementary Figure 1) are described in the Supplementary Information.

### Orbital Cultivation in the ISS

On the ISS, the culture vessels were set in the PEU and cultivated within CBEF, which has a rotating table for generating artificial gravity inside the Japanese Experiment Module “Kibo” (JAXA). In Run 1, cultivation was performed under 1G and µG conditions. In Run 2, cultivation was conducted under µG and 1/6G conditions.

Cultivation was controlled at 30 °C under a 14 h light / 10 h dark photoperiod, with the blue/red LED light intensity adjusted to a target of approximately 50 µmol photons m^-2^ s^-1^. Neutral density filters (ND 0.2; Fujifilm Corp., Tokyo, Japan) were used to compensate for LED output variations and ensure uniform irradiance across all experimental groups. Orbital cultivation was conducted for 8 days. During the experiment, the internal temperature and humidity of the PEU culture vessels were continuously monitored using Hygrochron™ iButton data loggers (Analog Devices, Inc., San Jose, CA, USA). Continuous monitoring confirmed that the internal temperature and humidity remained stable at approximately 29.7 °C and 96.1%, respectively, for both Run 1 and 2 throughout the cultivation period (Supplementary Figure 2).

### Image Processing and Quantification of Biomass Expansion

Images capturing the biomass expansion status of *Spirulina* were automatically acquired once daily using the internal RGB camera of the PEU and downlinked to the ground.

Prior to quantitative analysis, image alignment was conducted using the SIFT Feature Extraction plugin in Fiji (ImageJ) [11] to compensate for the spatial shifts of the PEU culture vessels over time. A specific region of interest (ROI) was manually defined along the boundary of the glass fiber filter that was not obstructed by the PEU culture vessel frame. Using an automated custom macro in Fiji, the green channel was extracted from the RGB images within the defined ROI to isolate the biomass, and the images were binarized using the automatic Moments thresholding method. To reduce image noise, morphological operations consisting of two cycles of dilation, followed by two cycles of erosion, were applied. Finally, the area fraction within the ROI was automatically measured. To evaluate radial growth on the solid substrate, the raw area fractions were converted to a relative radial growth metric calculated as √Area_Day X_ -√Area_Day 0_. The resulting values were used to plot the relative radial growth. To ensure analytical reproducibility, the image-processing workflow was semi-automated using a custom Fiji macro code generated and optimized with the assistance of Gemini 3.6 Flash (Google LLC, Mountain View, CA, USA). The complete code and processing parameters have been deposited in Zenodo (DOI: 10.5281/zenodo.23681047).

### Statistical analysis

Linear regression analysis was applied to the square root of the time-course biomass expansion data to determine relative radial growth rates (%^0.5^ day^-1^) based on the characteristically linear expansion profiles of attached microalgal cultures [12].

Goodness-of-fit was evaluated using the coefficient of determination (*R*^2^). To assess the effect of gravity while avoiding confounding factors, Student’s *t*-test (two-tailed) was performed independently within each experimental run: comparing 1G and µG in Run 1, 1/6G and µG in Run 2. Differences between the two µG controls were also evaluated using Student’s *t*-test. Statistical significance was set at *p* < 0.05. All statistical analyses were performed using Excel.

### Use of Generative AI and AI-assisted Technologies

During the preparation of this work the authors used Gemini in order to improve the English phrasing, correct grammatical errors, assist in structuring the manuscript draft, and consult on figure design and layout. After using this tool/service, the authors reviewed and edited the content as needed and take full responsibility for the content of the published article.

### Data Availability

The datasets generated and/or analyzed during the current study are available from the corresponding author on reasonable request.

### Code Availability

The custom Fiji macro code used for image processing and quantification of biomass expansion will be made publicly available in Zenodo upon publication of this article (DOI: 10.5281/zenodo.23681047).

## Acknowledgment

We sincerely thank ISS Science Laboratory Support Team at NASA Kennedy Space Center (KSC) for their support with the laboratory setup and logistics at KSC. We are deeply grateful to NG-24 mission team from Northrop Grumman for their coordination and efforts in preparing the payload onto the NG-24 cargo. We also extend our gratitude to the NASA Cold Stowage Team for their technical support in establishing the cold-chain transport for our samples. We are grateful to the ISS Expedition 74 crew members for their dedicated on-orbit operations, and the Japan Aerospace Exploration Agency (JAXA) Flight Control Team for their invaluable real-time operational support. This work was supported by the JAXA under the ’Kibo’ utilization program (Theme Name: Space Surface Spirulina, ID: 72749).

## Author Contributions

**T.I.** conceived the methodology, developed the software, performed formal analysis and data curation, conducted the investigation, visualized the results, wrote the original draft, and contributed to project administration, supervision, and manuscript editing. **M.K.** validated the data, conducted the investigation, managed resources and data curation, and edited the manuscript. **H.O.** conducted the investigation, contributed to supervision, and edited the manuscript. **T.S.** conducted the investigation, provided resources, and edited the manuscript. **J.K.** developed the methodology, managed project administration, and supervised the study. **Y.K.** conducted the investigation, managed resources and data curation, and edited the manuscript. **H.K.** conducted the investigation, performed data curation, and edited the manuscript. **T.Y.** developed the methodology and managed project administration. **K.K.** provided resources and edited the manuscript. **E.M.** contributed to study conceptualization, supervision, and manuscript editing. **S.A.** contributed to project administration and manuscript editing. **T.H.** conceptualized the study, secured funding, and edited the manuscript. All authors reviewed and approved the final version of the manuscript.

