## Supplementary Information for "A Proof-of-Concept Study on Attached *Spirulina* Cultivation Under Microgravity and Partial Gravity Aboard the ISS"

### Contents

- **Supplementary Method**
- **Supplementary Table 1.** Water contents of harvested biomass of attached *Spirulina* cultivation on the ground were approximately 90 wt%.
- **Supplementary Figure 1.** Pre-cultivation cold storage duration does not affect *Spirulina* biomass productivity within the 8-day timeframe of the spaceflight experiment.
- **Supplementary Figure 2.** Environmental conditions inside the PEU culture vessels throughout the entire spaceflight mission.
- **Supplementary Movie 1.** Time-lapse video showing the expansion of *Spirulina* biomass under 1G in Run 1.

- **Supplementary Movie 2.** Time-lapse video showing the expansion of *Spirulina* biomass under μG in Run 1.
- **Supplementary Movie 3.** Time-lapse video showing the expansion of *Spirulina* biomass under 1/6G in Run 2.
- **Supplementary Movie 4.** Time-lapse video showing the expansion of *Spirulina* biomass under μG in Run 2.

**Supplementary Method**

Approximately 15 mg of *Spirulina* was inoculated onto the surface of a 50 mm × 50 mm cellulose filter plate (No. 1034-3A, ADVANTEC TOYO, Tokyo, Japan) as a circular area with a diameter of 36 mm via suction filtration. Three sample-inoculated filter plates were placed side-by-side inside a square Petri dish (230 mm × 82 mm × 18 mm, AS ONE Corporation, Osaka, Japan), where a water-retentive sheet (220 mm × 75 mm, Germguard; Toyobo Co., Ltd., Osaka, Japan) and 45 mL of SOT medium were laid at the bottom. To prevent evaporation of the medium, the dish was enclosed in a polypropylene bag along with a moistened paper wipe (Kimwipe). The cultivation was conducted under continuous white LED light at 80 ± 10 μmol photons $m^{-2}$ $s^{-1}$ at 26–32 °C. Following cultivation, the biomass was harvested by scraping and weighed to determine the wet weight. The biomass was washed three times with Milli-Q water and dried overnight at 60 °C to evaluate biomass growth based on dry weight (Supplementary Figure 1) and calculate the moisture content (Supplementary Table 1).

**Supplementary Table 1. Water contents of harvested biomass of attached *Spirulina* cultivation on the ground were approximately 90 wt%.**

| Cold storage periods (weeks) | Cultivation time (day) | Wet biomass weight (mg) | Dry biomass weight (mg) | Water content (wt%) |
|---|---|---|---|---|
| 2 | 4 | 281.0 ± 6.4 | 24.9 ± 0.6 | 91.2 ± 0.1 |
| | 8 | 407.7 ± 50.0 | 42.6 ± 4.0 | 89.5 ± 0.4 |
| 4 | 4 | 308.0 ± 17.2 | 28.2 ± 1.2 | 90.8 ± 0.2 |
| | 8 | 372.4 ± 12.1 | 42.4 ± 3.5 | 88.6 ± 0.7 |

Values represent individual technical replicates ($n = 3$) and their mean ± standard deviation (SD). Moisture content was calculated using the formula: Moisture Content (wt%) = [(Wet Weight - Dry Weight)/Wet Weight] × 100

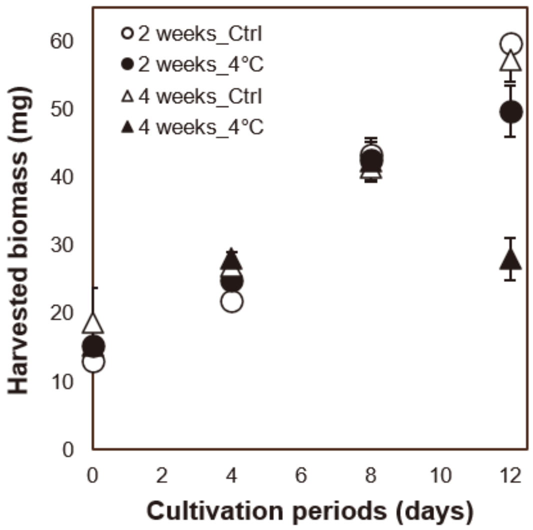


**Supplementary Figure 1. Pre-cultivation cold storage duration does not affect *Spirulina* biomass productivity within the 8-day timeframe of the spaceflight experiment.** Biomass productivity was measured over a 12-day cultivation period after cold storage. Values are presented as mean ± SD (n = 3, technical replicates).

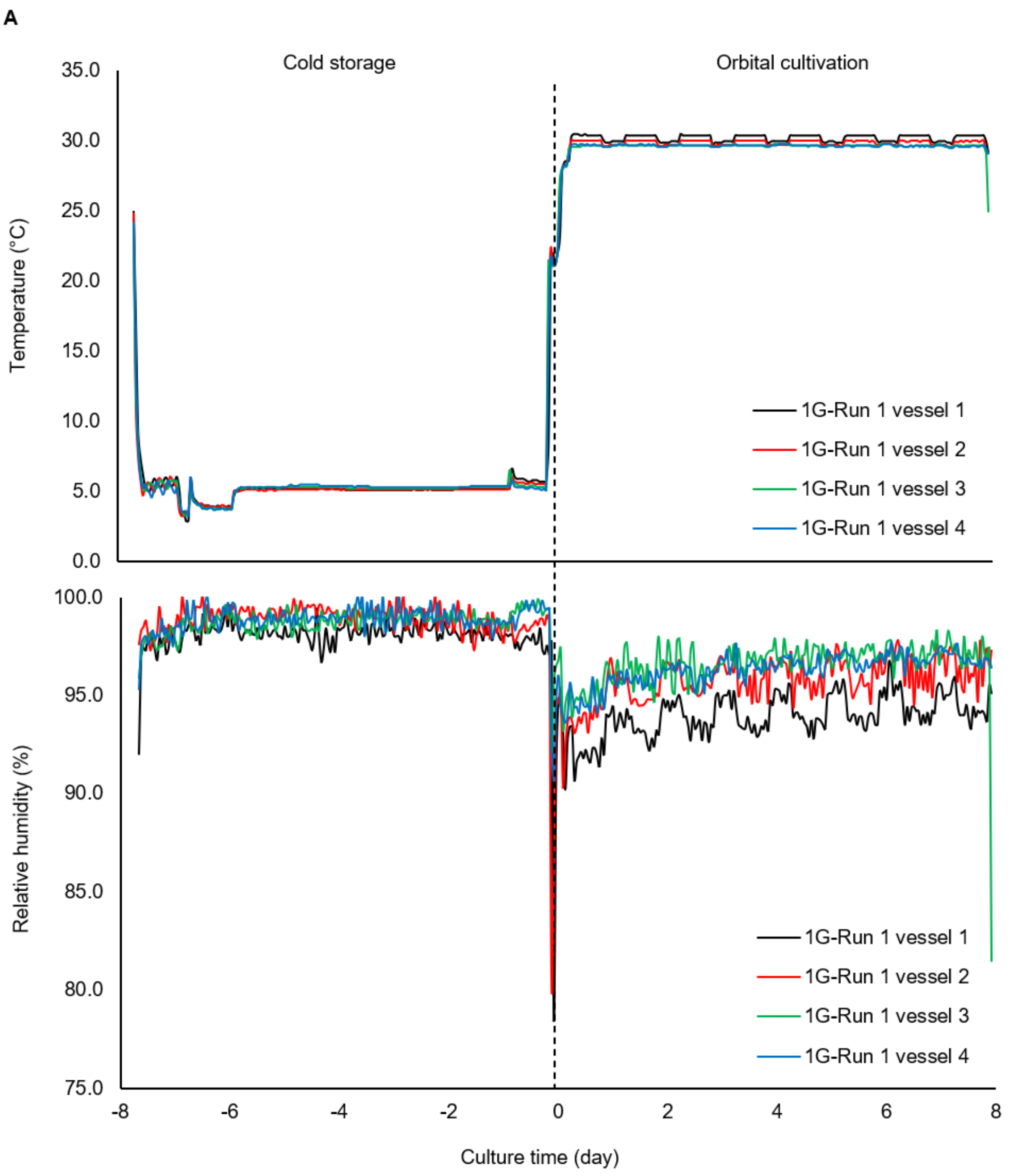


**Supplementary Figure 2. Environmental conditions inside the PEU culture vessels throughout the entire spaceflight mission.** Continuous temperature (top) and relative humidity (bottom) profiles under μG in Run 1 (A), 1G in Run 1 (B), 1/6G in Run 2 (C), and μG in Run 2 (D). These profiles were recorded by Hygrochron™ iButton data loggers during ground cold storage (prior to Day 0) and orbital cultivation aboard the

ISS (Day 0 to Day 8). Vertical dashed lines indicate the transition from cold stowage to the start of cultivation inside the CBEF.

Supplementary Figure 2 (Continued)

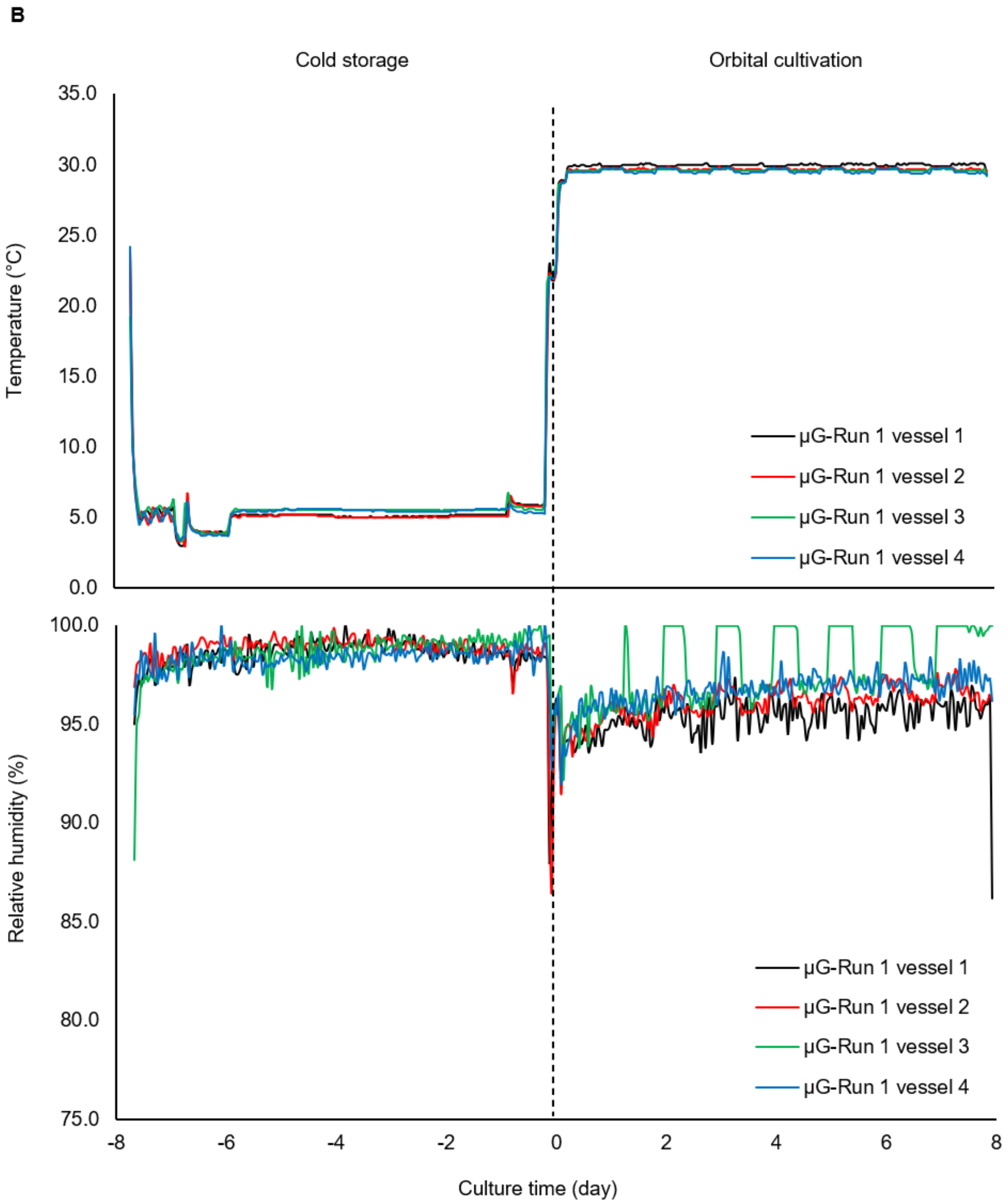

Supplementary Figure 2 (Continued)

c

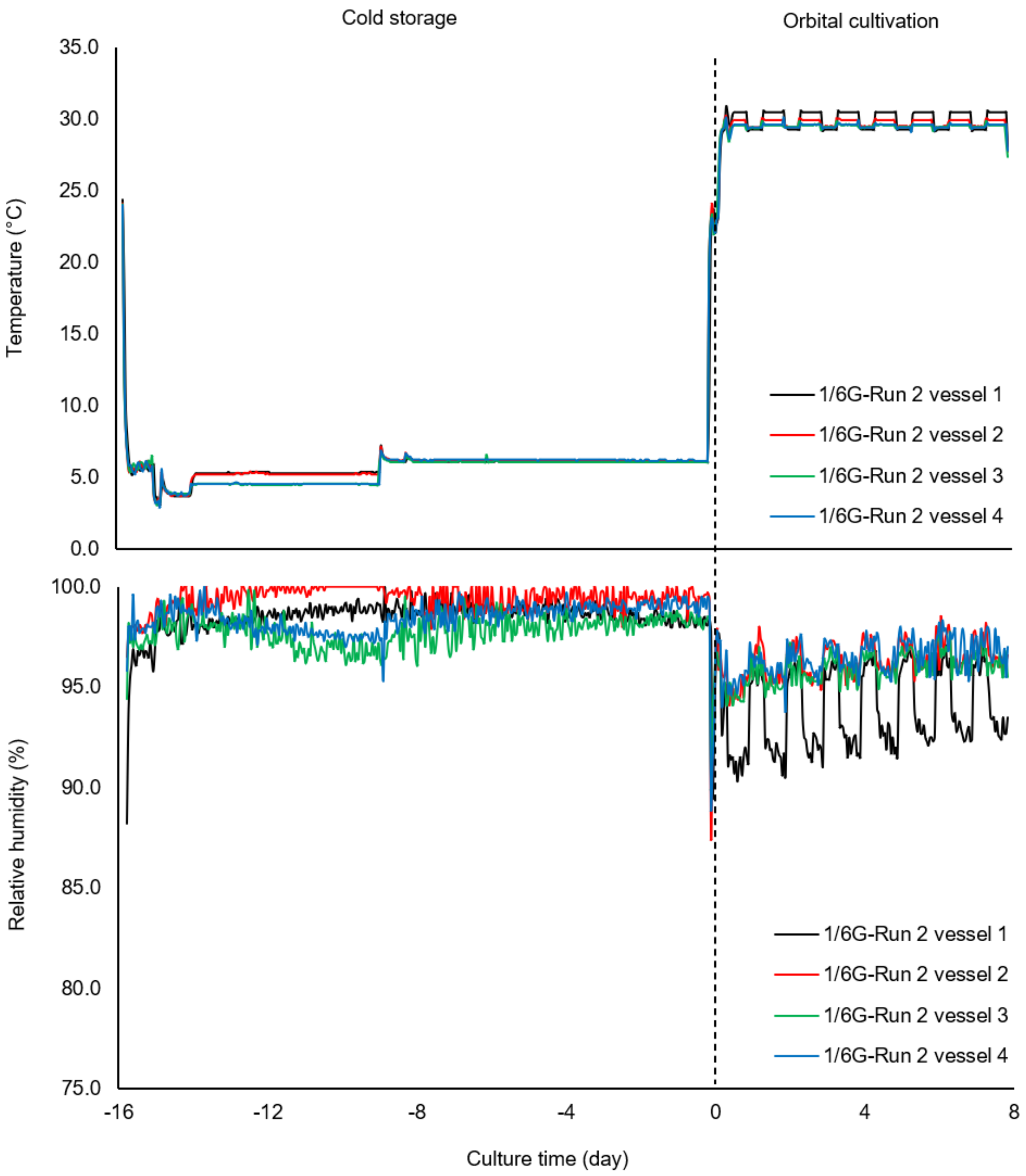

Supplementary Figure 2 (Continued)

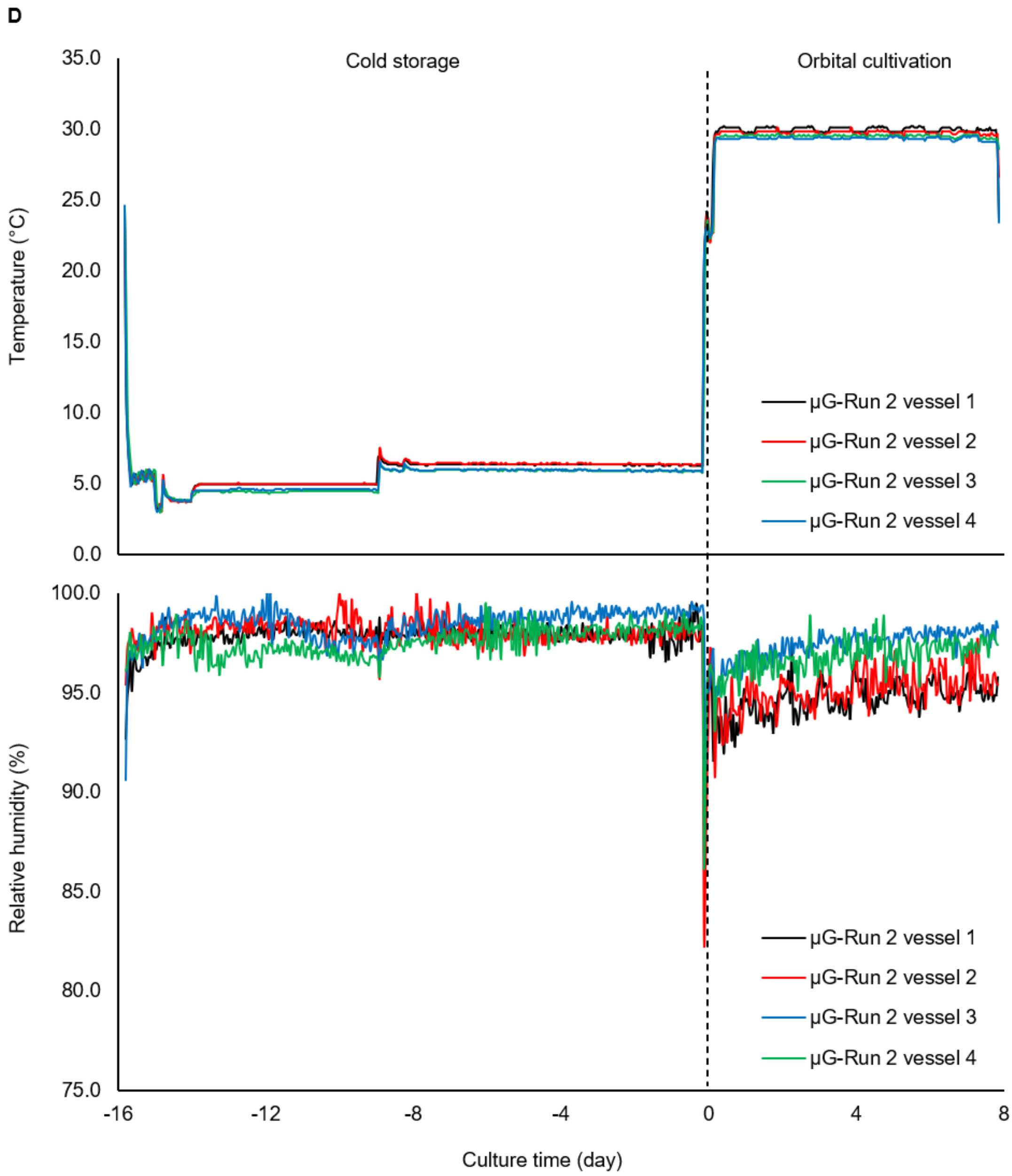

**Supplementary Movie 1. Time-lapse video showing the expansion of *Spirulina* biomass under 1G in Run 1.** Top panels show the aligned raw images and bottom panels show the binarized images from Day 0 to Day 8 ($n$ = 4).

**Supplementary Movie 2. Time-lapse video showing the expansion of *Spirulina* biomass under μG in Run 1.** Top panels show the aligned raw images and bottom panels show the binarized images from Day 0 to Day 8 ($n$ = 4).

**Supplementary Movie 3. Time-lapse video showing the expansion of *Spirulina* biomass under 1/6G in Run 2.** Top panels show the aligned raw images and bottom panels show the binarized images from Day 0 to Day 8 ($n$ = 4).

**Supplementary Movie 4. Time-lapse video showing the expansion of *Spirulina* biomass under μG in Run 2.** Top panels show the aligned raw images and bottom panels show the binarized images from Day 0 to Day 8 ($n$ = 4).
